# Genome-wide heritability analysis of severe malaria susceptibility and resistance reveals evidence of polygenic inheritance

**DOI:** 10.1101/649095

**Authors:** Delesa Damena, Emile R. Chimusa

## Abstract

**Objective:** Estimating SNP-heritability (*h^2^_g_*) of severe malaria/resistance and its distribution across the genome might shed new light in to the underlying biology.

**Method:** We investigated *h^2^_g_* of severe malaria susceptibility and resistance from genome-wide association study (GWAS) dataset (sample size =11, 657). We partitioned the *h^2^_g_* in to chromosomes, allele frequencies and annotations. We further examined none-cell type specific and cell type specific enrichments from GWAS-summary statistics.

**Results:** We estimated the *h*^2^*_g_* of severe malaria at 0.21 (se=0.05, p=2.7×10^−5^), 0.20 (se =0.05, p=7.5×10^−5^) and 0.17 (se =0.05, p= 7.2×10^−4^) in Gambian, Kenyan and Malawi populations, respectively. The *h^2^_g_* attributed to the GWAS significant SNPs and the well-known sickle cell (*HbS*) variant was approximately 0.07 and 0.03, respectively. We prepared African population reference panel and obtained comparable *h^2^_g_* estimate (0.21 (se = 0.02, p< 1×10^−5^)) from GWAS-summary statistics meta-analysed across the three populations. Partitioning analysis from raw genotype data showed significant enrichment of *h^2^_g_* in protein coding genic SNPs while summary statistics analysis suggests pattern of enrichment in multiple categories.

**Conclusion:** We report for the first time that the heritability of malaria susceptibility and resistance is largely ascribed by common SNPs and the causal variants are overrepresented in protein coding regions of the genome. Overall, our results suggest that malaria susceptibility and resistance is a polygenic trait. Further studies with larger sample sizes are needed to better understand the underpinning genetics of resistance and susceptibility to severe malaria.

## INTRODUCTION

Malaria remains one of the leading public health problems worldwide. The global tally of malaria in 2017 was estimated at 219 million cases and 435 000 deaths worldwide (1). *P. falciparum* malaria results in diverse clinical manifestations ranging from asymptomatic parasitaemia to severe malaria (2,3). Family studies reported that the host genetic factors (heritability) contributes about 25% of the variations observed in clinical severity of malaria (4). Heritability (H^2^) is defined as the proportion of the phenotypic variations attributable to genetic differences in a population (5). Narrow-sense heritability (*h*^2^) is a specific term that refers to the phenotypic variations attributable to the additive genetic values. Quantifying the proportion of narrow-sense heritability attributed to common variants from GWAS SNPs without identifying the causal variants, commonly called SNP-heritability (*h^2^_g_*), has recently gained a considerable interest in statistical genetics to address the problem of ‘missing’ heritability in GWASs (6).

Several statistical methods now allow accurate estimation of *h^2^_g_* from all SNPs of unrelated individuals (7–9). Methods such as the Genome-wide Complex Trait Analysis (GCTA) compute genetic relationships matrix (GRMs) of unrelated subjects by simultaneously fitting all SNPs and estimates the proportion of phenotypic variations explained by genotypic variation using restricted maximum likelihood (GREML) methods (7). Phenotype-Correlation-Genotype-Correlation (PCGC) is a one of recently developed methods specifically designed to estimate *h^2^_g_* from case/control (9). PCGC is a Haseman-Elston (H-E) regression model in which normalized phenotype is regressed on the genetic covariances of all unique pair of samples. The slope of the regression is used as an estimator of SNP-heritability. Application of these methods provided new insights in to the genetic architecture of complex disease genetic architecture (10, 11). Alternative contemporary statistical methods that enable estimation of *h^2^_g_* from publicly available GWAS-summary statistics without the need of individual genotype data are also widely available (12–14) and gained popularity due to their privacy advantages and computational costs.

Even though a number of severe malaria GWASs have recently been implemented in malaria endemic areas in Africa and reported a number novel variants (15–18), little is known about the proportion of *h^2^_g_* and its distribution across chromosomes, functional regions and allele frequency spectrum. We hypothesize that malaria susceptibility and resistance is a complex trait that is mainly under polygenic control. Here we present results from comprehensive SNP-heritability studies of severe malaria in three African populations including Gambia, Kenya and Malawi. We estimated *h^2^_g_* of severe malaria and partitioned in to chromosomes, different minor allele frequency (MAF) bins annotations and cell-types. Overall, our results suggest that malaria susceptibility and resistance is a polygenic trait.

## RESULTS

### SNP-heritability of severe malaria susceptibility and resistance from genotype data

We estimated SNP-heritability at different quality control (QC) levels to determine the appropriate threshold (see Materials and Methods). As expected, the estimates were inflated when less stringent QC thresholds were applied (**Supplementary Figure 1**). However, when more stringent sets of QCs including relatedness threshold (5%), SNP differential missingness proportion (p < 1× 10^−3^) and SNPs missing proportion (p > 0.02)) were applied, the estimates became stable in each population (Gambia, Kenya and Malawi). Inclusion of more PCs (15, 20, 50) as a fixed effect didn’t further decrease these estimates. At the stringent QC threshold, the SNP-heritability of severe malaria was 0.21 (se=0.05, p=2.7×10^−5^), 0.20 (se =0.05, p=7.5×10^−5^) and 0.17 (se =0.05, p= 7.2×10^−4^) in Gambian, Kenyan and Malawi populations, respectively (**Table 1**). The SNP-heritability was approximately similar in major ethnic groups including Mandinka (0.24, se=0.06, p=5.1×10^−5^) in Gambia and Chonye (0.20, se = 0.07, p = 5.1× 10^−3^) and Girimia (0.19, se=0.07, p=7.3× 10^−3^) in Kenya. We didn’t estimate *h^2^_g_* for other ethnic groups because of smaller sample sizes. Furthermore, the PCGC model approximately showed similar results: 0.20, (se=0.06, p=9.7×10^−4^), 0.16 (se=0.06, p=8×10^−3^) and 0.23 (se=0.07, p=1.3×10^−3^) in Gambia, Kenya and Malawi populations, respectively.

**Table 1.**
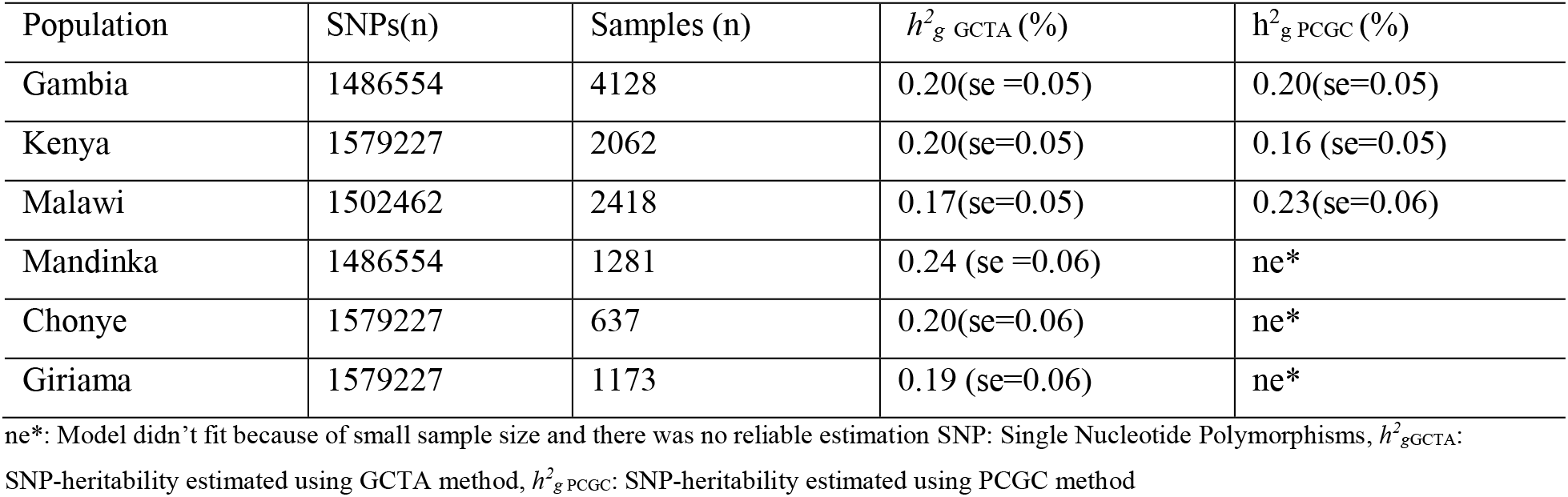
SNP-heritability of severe malaria susceptibility and resistance determined by GCTA and PCGC methods.

| Population | SNPs(n) | Samples (n) | $h^2_g$ GCTA (%) | $h^2_g$ PCGC (%) |
| --- | --- | --- | --- | --- |
| Gambia | 1486554 | 4128 | 0.20(se =0.05) | 0.20(se=0.05) |
| Kenya | 1579227 | 2062 | 0.20(se=0.05) | 0.16 (se=0.05) |
| Malawi | 1502462 | 2418 | 0.17(se=0.05) | 0.23(se=0.06) |
| Mandinka | 1486554 | 1281 | 0.24 (se =0.06) | ne* |
| Chonye | 1579227 | 637 | 0.20(se=0.06) | ne* |
| Girima | 1579227 | 1173 | 0.19 (se=0.06) | ne* |

ne\*: Model didn't fit because of small sample size and there was no reliable estimation SNP: Single Nucleotide Polymorphisms, $h^2_{gGCTA}$ :
SNP-heritability estimated using GCTA method, $h^2_{gPCGC}$ : SNP-heritability estimated using PCGC method

### Heritability explained by GWAS significant SNPs

To quantify the effects of the known variants, we estimated *h^2^_g_* without removing malaria associated loci (see Materials and Methods). This resulted in slight increase of the estimate in Gambian (0.27 (se = 0.0 5, p = 1×10^−5^)) and Kenyan (0.24 (se = 0.05, p < 1×10^−5^)) populations. Including *rs334* as an additional covariate decreased the estimate to (0.24 (se = 0.05, p < 1×10^−5^)), 0.21 (se =5%, p < 1×10^−5^)) in Gambian and Kenyan populations, respectively. Suggesting that the *h^2^_g_* attributed to the significant SNPs and *HbS* locus is about 0.07 and 0.03, respectively across the populations.

### Partitioning SNP-heritability by chromosomes and minor allele frequencies

We observed no significant differences in between the *h^2^_g_* per chromosome obtained from separate analysis (0.24, se = 0.05, p < 1×10^−5^) and that obtained from the joint analysis (0.20, se = 0.05, p = 1×10^−5^) as shown in **Supplementary Figure 2**, suggesting that the population structure was adequately controlled. Moreover, we observed significant correlations between chromosomal length and *h^2^_g_* per chromosome (Adj r^2^ = 0.38, p = 0.001) as shown in **Figure 1**. However, the estimates of three chromosomes (chr 5, 11 and 20) and three other chromosomes (chr 7, 8 and 15) fell above and below the expected *h^2^_g_* at 95% CI, respectively. Notably, chr 5 contain a considerable proportion (15%) of the cumulative *h^2^_g_* estimate. To determine the relative contribution of variants from various allele frequencies, we partitioned SNPs in to different MAF bins and estimated *h^2^_g_* attributed to each bin using joint GRMEL analysis (**Figure 2**). However, we didn’t detect significant differences between the proportion of *h^2^_g_* attributed to different MAF bins (standard errors overlapped at 95% CI). The total sum of *h^2^_g_* across bins (0.27 (se = 0.08, p = 5.3×10^−5^)) was not significantly different from the univariate estimate.

**Figure 1.**
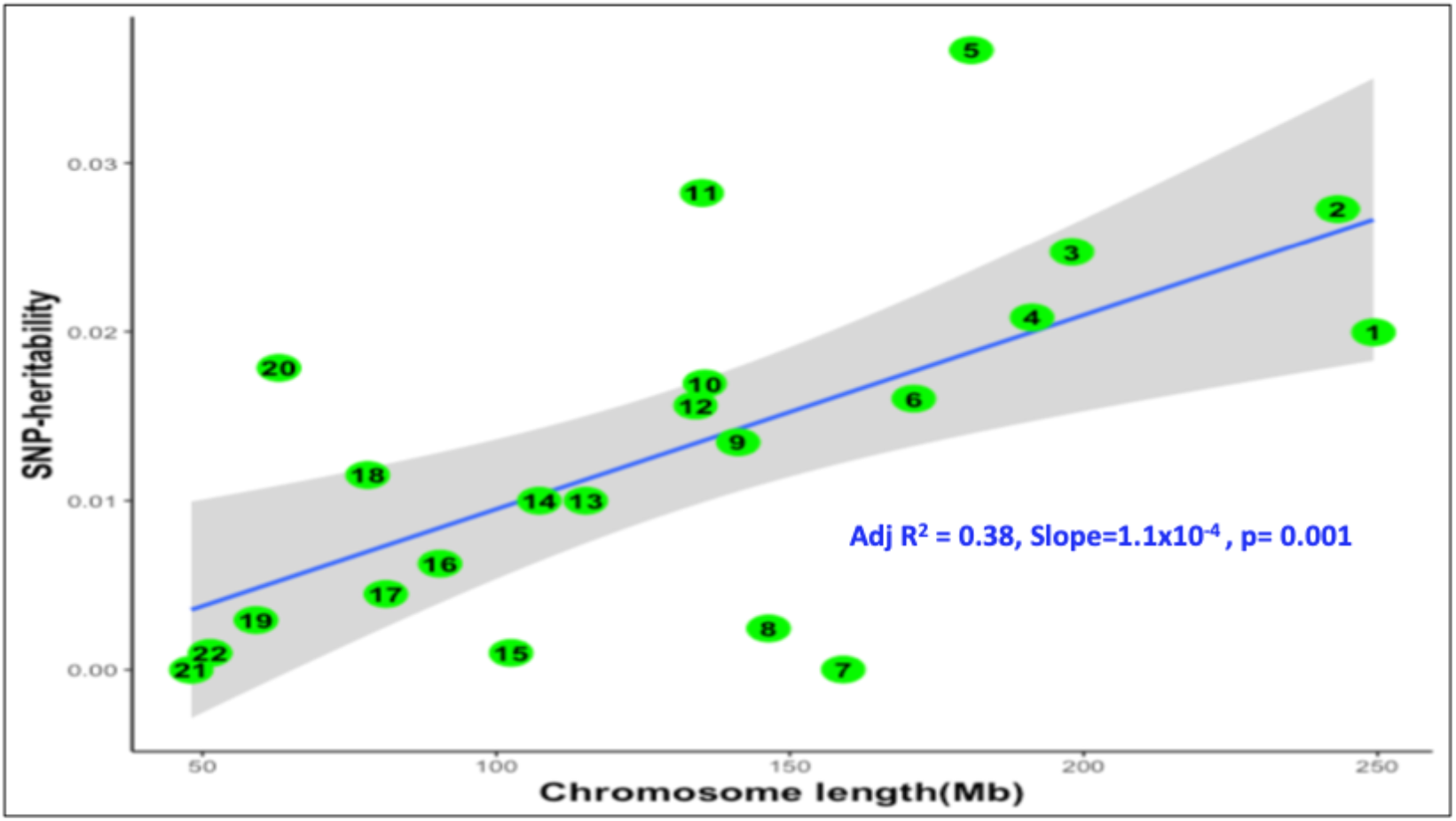
SNP-heritability (*h^2^_g_*) per chromosome(y-axis) plotted against chromosome length (x-axis). The blue line represents the *h^2^_g_* estimates regressed against chromosome length. The grey shaded areas represent the 95% confidence interval around the slope of the regression model.

**Figure 2.**
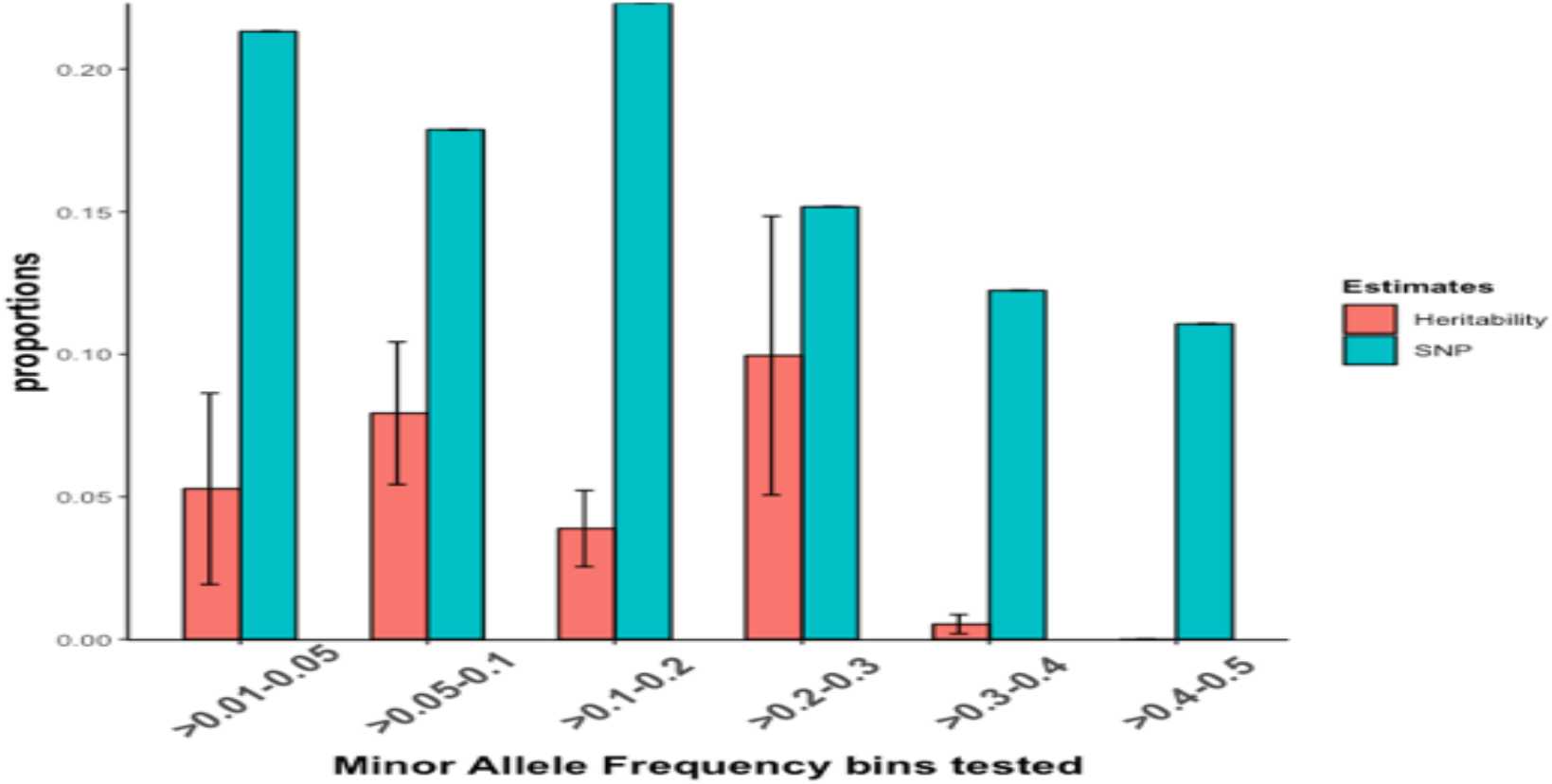
SNP-heritability partitioned in to different allele frequency spectrum. We created six MAF-bins and estimated the proportion of *h^2^_g_* attributed to each bin. The proportion of *h^2^_g_* attributed to each bin is shown in red bar and the proportion of SNPs per bin is shown by blue bar. Error bars represent the 95% CI of the estimate.

### Partitioning SNP-heritability by annotation

We estimated the *h^2^_g_* explained by genic SNPs and intergenic SNPs at 0.163 (se=0.06) and 0.064 (se=0.05), respectively (**Table 2**). This implies that the proportion of heritability explained by a SNP residing in a gene within 10kb boundary is on average twice as the heritability explained by a SNP residing in an intergenic region. However, the statistical inferences can’t be made as standard errors overlapped at 95% CI. We further calculated the *h^2^_g_* explained by protein coding genic SNPs and the corresponding intergenic SNPs at 0.165 (se = 0.05) and 0.062 (se = 0.05), respectively. A SNP residing in protein coding genes within 10kb boundary contribute about 2.9 x *h^2^_g_* than a SNP residing in an intergenic region and this is statistically significant at 95% confidence interval.

**Table 2.**
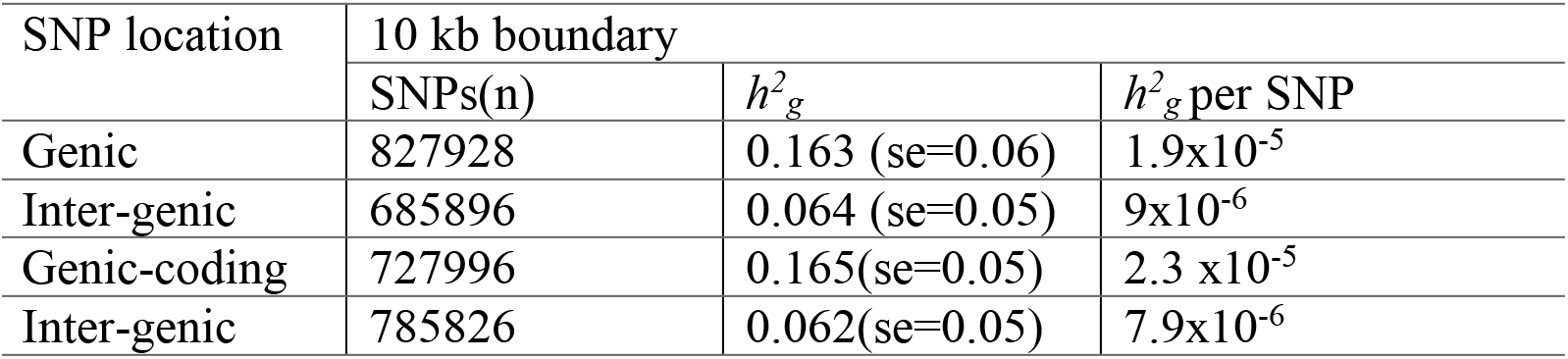
SNP-heritability of severe malaria susceptibility and resistance partitioned in to genic and intergenic genomic regions.

### Functional enrichment from GWAS-summary statistics

After imputation from malaria GWAS-summary statistics and QC filtering, we obtained a total of 20 million high quality SNPs (see Materials and Methods). Using this dataset, we estimated the liability scale SNP-heritability at 0.21 (se = 0.02, p < 1×10^−5^). Partitioning the *h^2^_g_* in to 24 main genomic annotations (baseline model) showed patterns of enrichment in multiple categories including 5’UTR (11×), DGF (10x), enhancer (9x), coding (6x), H3K4me1 (4.9x), TSS (5x), TFBS (4×), and FANTOM enhancer (4×) as shown in **Figure 3**. However, none of the enrichments was statistically significant after correction for multiple testing. Further cell-type specific and cell group analysis didn’t show significant enrichment.

**Figure 3.**
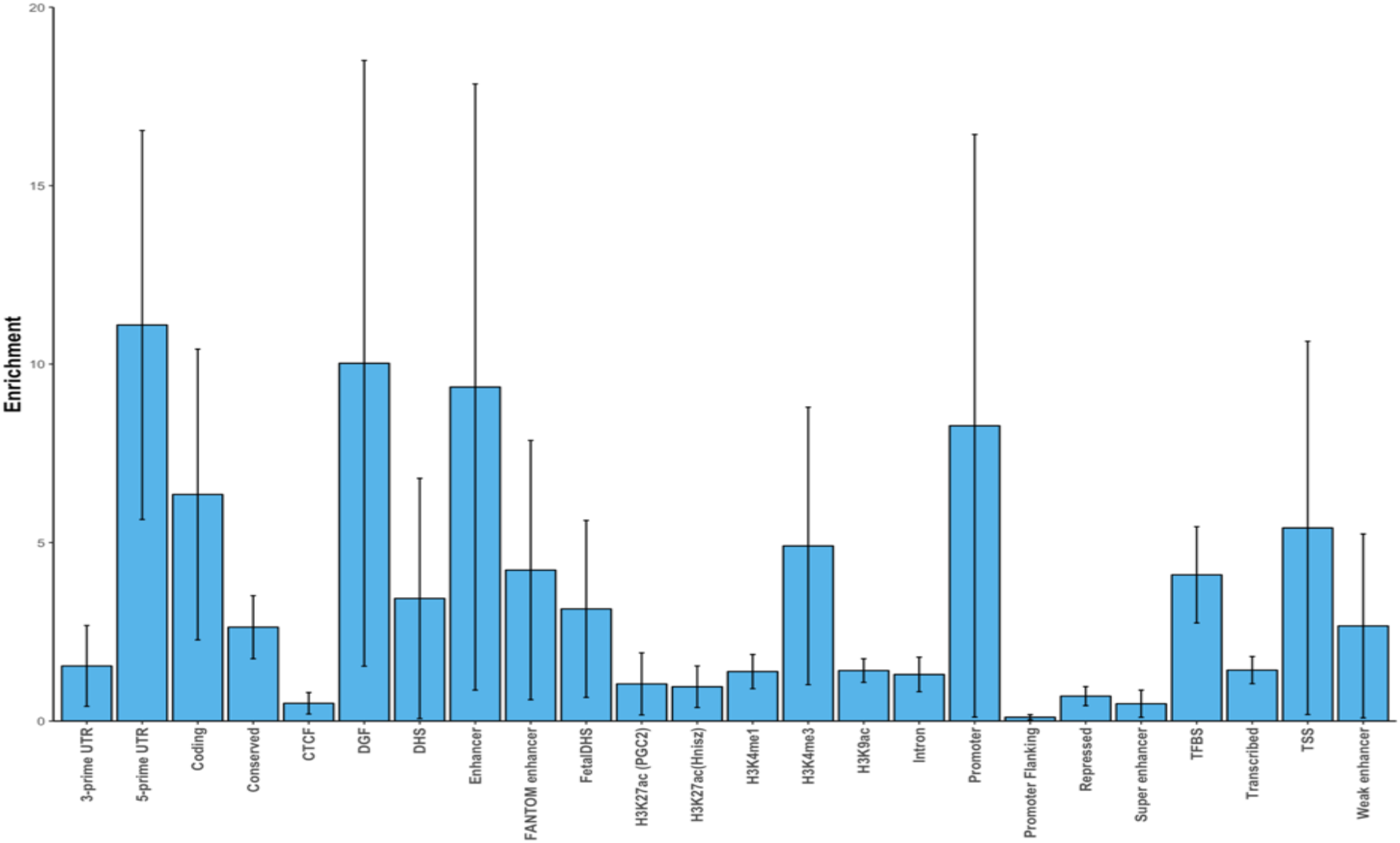
Enrichment estimates of severe malaria susceptibility and resistance for the 24 main annotations. Error bars represent jackknife standard errors around the estimates of enrichment

## DISCUSSIONS

In this study, we estimated the SNP-heritability and functional enrichment of malaria susceptibility and resistance in three African populations and their meta-analysis. We performed GRMEL analysis at different QC levels to determine the appropriate threshold; indeed, the estimates were inflated upward at less stringent QC levels and became stable at more stringent QC levels. At the stringent QC threshold, the *h^2^_g_* of severe malaria susceptibility was 0.21 (se=0.05, p=2.7×10^−5^), 0.20 (se =0.05, p=7.5×10^−5^) and 0.17 (se = 0. 05, p = 7.2×10^−4^) in Gambian, Kenyan and Malawi populations, respectively. The estimates were approximately similar across major ethnic groups in the study populations. Application of PCGC method in our dataset yielded comparable estimates including 0.20, (se=0.06, p=9.7×10^−4^), 0.16 (se = 0.06, p = 8×10^−3^) and 0.23 (se =0.07, p=1.3×10^−3^) in Gambia, Kenya and Malawi populations, respectively.

Despite slight differences, our *h^2^_g_* estimate was approximately the same in different populations, arguing that the allele heterogeneity of the malaria susceptibility/resistance and their effect sizes (3) might not substantially affect the overall SNP-heritability across endemic populations. Replication studies with larger population specific sample sizes might be needed to further investigate this observation.

Our *h^2^_g_* estimates was roughly close to a previous report from a family-based study in which the narrow-sense heritability estimates of severe malaria was estimated to be 0.25 (4). The same study estimated the contribution of the well-known *HbS* variant at 0.025 which is also consistent with our estimate (0.03). Suggesting that the malaria susceptibility/resistance is influenced by several variants each individually having a small effect sizes in population and thus, it is a polygenic trait. This is consistent with the hypothesis that the vast proportion of heritability of complex traits is attributed to SNPs with effect sizes too small to attain the stringent genome wide significance threshold (p < 5 x 10^−8^) at the current sample sizes (19). To gain better insights in to the genetic architecture of severe malaria, we partitioned the *h^2^_g_* in to different chromosomes, allele frequency spectrum and annotations using joint analysis implemented in GCTA software (7). We found no significant difference between the separate analysis (0.25, se = 0.05, p < 1×10^−5^) and joint analysis (0.24, se=0.05, p =1×10^−5^), suggesting that the population structure is adequately controlled. The rationale is that separate analysis in which one chromosome is fitted to the model at a time, captures variances due to correlated SNPs on other chromosomes and thus, yield inflated estimates in the presence of population structure. However, the joint analysis in which GRMs of all chromosomes are simultaneously fitted in to a single GRMEL, effectively controls upward biases that can be created from correlated SNPs on different chromosomes and thus, provide more accurate estimates (29).

Supporting the polygenic view of genetic architecture, we found a correlation between *h^2^_g_* per chromosome and chromosomal lengths (Adj r^2^ = 0.39, p = 0.001). However, the estimates of three chromosomes (chr 5, 11 and 20) and three other chromosomes (chr 7, 8 and 15) fell below above and below the expected *h^2^_g_* at 95% CI. Notably, chr 5 contain a considerable proportion 15% (0.036) of the cumulative *h^2^_g_* estimate, suggesting that this chromosome might contain loci with larger effects against the polygenic background. Previous family-based studies reported the association of 5q31-q33 with mild malaria susceptibility and resistance (20, 21).

MAF-stratified analysis didn’t reveal significant differences between the proportion of *h^2^_g_* attributed to different MAF bins (standard errors overlapped at 95% CI). This might assert that *h^2^_g_* of severe malaria is broadly uniform across the allele frequency spectrum and is not over-represented by rare alleles. Moreover, the total sum of *h^2^_g_* in MAF-stratified analysis (0.27 (se = 0.08, p = 5.3× 10^−5^)) was not significantly different from the cumulative univariate *h^2^_g_* estimate, further suggesting that allelic heterogeneity of malaria protective variants in different populations might not have substantial effects on the overall SNP-heritability. However, more powered studies might be needed to replicate this observation.

Partitioning by annotation revealed that the *h^2^_g_* of severe malaria susceptibility and resistance is concentrated in genic regions of the genome. The proportion of heritability explained by a SNP in genes within 10kb boundary is on average twice as the heritability explained by a SNP in intergenic regions. On average, a SNP in protein coding genes within 10kb boundary contribute about 2.9 x *h^2^_g_* than a SNP in intergenic regions. The later was statistically significant at 95% CI. This finding is arguing against the notion that SNP-heritability of complex traits is largely attributed to SNPs residing in none coding regions of the genome.

In addition to the direct estimation from the raw genotype, functional enrichment analysis from GWAS-summary statistics using stratified LDSC (12) method has recently been shedding new lights in to complex disease genetic architectures (22). However, this approach require population specific reference panel to produce reliable results making it difficult to apply in African populations which has been poorly represented in public reference domain (12). In effort to address this challenge, we prepared African population specific reference panel for the first time which can be obtained from http://web.cbio.uct.ac.za/~emile/software.html. Our liability scale SNP-heritability estimate from GWAS-summary statistics meta analysed across the study populations (0.21 (se = 0.02, p < 1×10^−5^)) was comparable to the direct estimates from raw genotype data. However, our full base line model and cell specific didn’t reveal significant enrichments. Of note, the coding and regions surround coding genes were among the top categories in our base line model, further implying the importance of genic regions of the genome in influencing the malaria susceptibility and resistance trait. One of the downsides of stratified LDSC analysis is that it requires very large sample sizes to detect significant enrichments. We strongly believe that replication studies with large sample sizes can shed new lights in to the genetic architecture of severe malaria susceptibility and resistance.

## CONCLUSIONS

In conclusion, our study showed for the first time that heritability of malaria susceptibility and resistance is largely explained by common SNPs and is disproportionately enriched in protein coding regions of the genome. Consistent with polygenic genetic architecture, we observed broadly even distribution of SNP-heritability across chromosomes and different allele frequency spectrum, implying that very large sample sizes will be needed to identify novel variants. We prepared African population specific reference panel and showed that LDSC analysis can provide reliable SNP-heritability estimates in African populations. Further studies with larger sample sizes are needed to understand the unpinning genetic architecture of severe malaria susceptibility/resistance phenotype.

## METHODS

### Description of the study datasets

GWAS data of three African populations including Gambia, Kenya and Malawi were obtained from European Phenome Genome Archive (EGA) through the MalariaGen consortium standard data access protocols (23, 24). The dataset contains information about a total of 11, 657 samples including 4921 samples from Gambia (2491 cases and 2430 controls), 3752 samples from Malawi (3752 cases and 13220 controls), 2984 samples from Kenya (1506 cases and 1478 controls). Cases were obtained from children who were admitted to Hospitals and fulfilled WHO case definition for severe malaria (24) and controls were obtained from the general population (15–18). All the samples were genotyped on Illumina Omni 2.5M array and were provided in VCF format. Information about phenotypes, imputation and quality control (QC) was provided with the data.

### Quality control

The basic QC analyses including plate effects, sample relatedness, Hardy-Weinberg equilibrium, heterozygosity, missingness and differential missingness were conducted for the previously published GWASs as described elsewhere (15, 25). Taking in to consideration that small artifacts can have substantial cumulative effects in heritability analysis (26), we applied further stringent QC protocols. Briefly, we aligned the quality filtered VCF files to the forward strand of the human reference sequence (GRCh37), using the Illumina-supplied files (www.well.ox.ac.uk/~wrayner/strand) and removed all SNPs with position and strand mismatches. We further removed SNPs with minor allele frequency (MAF) below 0.01, deviate from Hardy-Weinberg at p-value below 0.01 using PLINK software (27). We then implemented step-wise QC control based on SNPs missingness proportion, differential missingness and sample relatedness.

### Estimating heritability from genotype data

We applied GCTA (7) to estimate the SNP-heritability of severe malaria susceptibility and resistance from raw genotype datasets. Briefly, we excluded the region of extended inversion (723,8552-12,442,658) on chromosome 8p23 (28), the Major Histocompatibility Complex (MCH) region (25,000,000-40,000,000) on chr6 and the known malaria susceptibility associated loci including the *ATP2B4* region on chr1:203,154,024-204,154,024, cluster of *glycophorin (GYPA/B/E)* region on chr4:143,000,000-146,000,000, *ABO* blood group region on chr9:135,630,000-136,630,000, and the sickle cell *(HbS)* region on chr11:2,500,000-6,500,000 to avoid potential biases.

We constructed Genomic Relatedness Matrix (GRMs) from pruned high quality independent autosomal SNPs (independent pairwise 50 10.2) using GCTA software (7) and obtained list of samples with relatedness threshold above 5%. We then computed GRMs using all SNPs for each cohort and excluded one of any pair of samples with relatedness threshold above 5% as recommended elsewhere (8). The final sample of unrelated individuals was 4128, 2062 and 2418 for Gambia, Kenya and Malawi, respectively. The distribution of off-diagonal element of the GRMs for each population is shown in **supplementary Figure 3**.

We estimated SNP-heritability (*h^2^_g_*) using GCTA software (7). We performed principal components analysis (PCA) and included the top 10 PCs as fixed effects to account for population structure. We used population prevalence of 1% of severe malaria as previously described in (24). We transformed the estimates to liability scale as outlined in Lee *et al.* (29). Using the same GRMs, we estimated the *h^2^_g_* using PCGC model as outlined in Golan *et al.* (12). We also computed separate GRMs for ethnic groups including Mandinka of Gambia and, Girimia and Chonye of Kenya and estimated the *h^2^_g_* attributed to each ethnic group. To quantify the effects of significant SNPs, we created GRMs without excluding the association loci and performed GRMEL analysis. We repeated the analysis by including rs334 as additional covariate to estimate the *h^2^_g_* attributed to *HbS*.

### Partitioning SNP-heritability from genotype data

Using Gambian dataset (largest sample size), we partitioned *h^2^_g_* by chromosomes, MAF bins and annotations. To investigate the biases that might be created by population structure, we computed GRMs for individual chromosome and estimated *h^2^_g_* attributed to each chromosome by separate GREML in which one chromosome is fitted to the model at a time. We then performed a joint analysis in which GRMs of all chromosomes are simultaneously fitted in to a single GREML analysis and compared the results obtained from both analyses. We further created five MAF bins including >0.01-0.05, >0.05-0.1, >0.1-0.2, >0.2-0.3, >0.3-04, >0.4-0.5, constructed separate GRMs for each bin and estimated *h^2^_g_* attributed to each category using the joint analysis implemented in GCTA software (7). Finally, we partitioned the SNP-heritability in to genic and intergenic regions of the genome. Briefly, we mapped all the autosomal SNPs to the reference genome using QCTOOLV2 (https://www.well.ox.ac.uk/~gav/qctool). We labelled all SNPs that are mapped to genes as genic and the remaining SNPs with no annotation as intergenic. We defined gene boundaries as regions within 10 kilobases (kb) regions upstream and downstream of a gene based on UCSC Genome Browser hg19 assembly annotation databases (http://genome.ucsc.edu). We estimated the *h^2^_g_* explained by the respective category using joint analysis in GCTA software (7).

### Functional enrichment analysis of SNP-heritability from GWAS-summary statistics

#### Preparation of African-specific reference panel

Partitioning *h^2^_g_* in to cell-types and functional categories using stratified Linkage Disequilibrium score regression (LDSC) approach has recently been shedding new lights in to the genetic architecture of several complex diseases (12, 22, 30). The method is based on the fact that a given category of SNPs is enriched for *h^2^_g_* if SNPs with high LD to that category have higher *χ*^2^ statistics than SNPs with low LD to that category (22, 30). However, the LDSC analysis require population specific reference panel and very large sample sizes to produce reliable results (12). Consequently, the current European 1000G haplotype reference panel that is used as a default in LDSC software(12, 30) might not well-represent our study populations. To address this challenge, we first merged African population datasets obtained from 1000 Genomes Project and African Genome Variation Project (31) based on overlapped variants and removed structural variants and ambiguous SNPs using plink tool (27). This resulted in a merged dataset of sample size (n = 4975). From this dataset, we excluded admixed populations including Americans of African Ancestry and African Caribbean. By taking in to consideration the high genetic diversity of African populations, we clustered the dataset into sub-regions comprised of eastern and western Africans using smart pca software (32) as shown in **Supplementary Figure 4**. We performed QC filtering based on MAF < 1%, missingness test (GENO > 0.05) and HWE test in controls (alpha level 0.0001). We retained a total of 22,473,268 SNPs and 18,919,068 SNPs SNPs in eastern (Sample size = 2,112) and western (Sample size = 380) African sub-region datasets. We eventually calculated the MAF of the panel for later partitioning analysis. Owing to the fact that our study populations are comprised of both east African (Malawi and Kenya) and west Africa (Gambian) populations, we utilized our entire dataset as a reference panel.

#### Baseline model and functional annotations

We created baseline model and cell type specific annotations for our reference panel as described in (12). The baseline-LD model included 24 main annotations that are not cell-type specific including coding, UTRs (3’UTR and 5’UTR), promoter and intronic regions obtained from UCSC genome browser and processed by Gusev *et al.* (10), the histone marks (H3) such as: acetylation of histone at lysine 9 (H3K9ac), monomethylation (H3K4me1) and trimethylation (H3K4me3) of H3 at lysine 4 obtained from Trynka *et al.* (33), acetylation of H3 at lysine 27(H3K27ac) version one processed by Hnisz *et al.* (34) and version two Psychiatric Genomics Consortium (PGC), combined chromHMM and Segway predictions obtained from Hoffman *et al.* (35), regions that are conserved in mammals (36, 37), super enhancers (34), FANTOM5 enhancers (38), transcription factor binding sites (TFBS) and digital genomic footprint (DGF) post-processed by Gusev *et al* (10). Around each category, we added 500bp windows as separate categories to prevent biases that might arise from adjacent categories. The 24 main annotations together with the additional windows and a category containing all SNPs yielded 53 overlapping baseline model. Next, we created 220 cell-type-specific annotations for the four histone marks: H3K4me1, H3K4me3, H3K9ac, and H3K27ac (12) using our reference panel and computed LD score for each annotation. We then combined the 120 cell specific annotations in to 10 cell groups including adrenal and pancreas, central nervous system (CNS), cardiovascular, connective and bone, gastrointestinal, immune and hematopoietic, kidney, liver, skeletal muscle, and other as described in (12). For each of the 10 categories, we computed the corresponding LD scores.

#### Application LDSC analysis to malaria susceptibility GWAS-summary statistics

We obtained meta-analysed GWAS-summary statistics of the three populations (n =15122094 SNPs) from the previous GWAS (15). We performed imputation on this dataset using ImpG software (39). Briefly, we removed SNPs that mismatch with 1000G phase three markers, computed z-score from the association statistics and performed the imputation using ImGv.1.1 under default settings. We used all 661 individuals labelled as “AFRICAN” haplotypes in phase 1 of 1000 Genome Project version-3 calls (40). We removed all imputed SNPs with a predicated accuracy less than 0.9 and SNPs with MAF < 0.01. After QC filtering, we performed stratified LD score regression analysis using our reference panels as described in (12). Briefly, we converted the summary statistics to LDSC format, filtered SNPs with imputation accuracy greater than nine and MAF greater than 1%, removed structural variants, ambiguous SNPs, the MHC region and significant SNPs. We then performed none-cell type and cell type specific analyses as described in (12).

## ACKNOWLEDGEMENTS

We thank Kwiatkowski’s group from University of Oxford for their constructive comments and assistance. We thank Gavin Band for his supervision and guidance in heritability analysis from genotype data. This work was supported through the DELTAS Africa Initiative [grant 107740/Z/15/Z]. The DELTAS Africa Initiative is an independent funding scheme of the African Academy of Sciences (AAS)’s Alliance for Accelerating Excellence in Science in Africa (AESA) and supported by the New Partnership for Africa’s Development Planning and Coordinating Agency (NEPAD Agency) with funding from the Wellcome Trust [grant 107740/Z/15/Z] and the UK government. The views expressed in this publication are those of the author(s) and not necessarily those of AAS, NEPAD Agency, Wellcome Trust or the UK government.

## AUTHORS CONTRIBUTIONS

DD designed, performed the data analysis and drafted the manuscript, EC contributed in designing, data-analysis and revision of the manuscript and supervised the work.

