## Supplementary Materials for "Genome-wide heritability analysis of severe malaria susceptibility and resistance reveals evidence of polygenic inheritance"

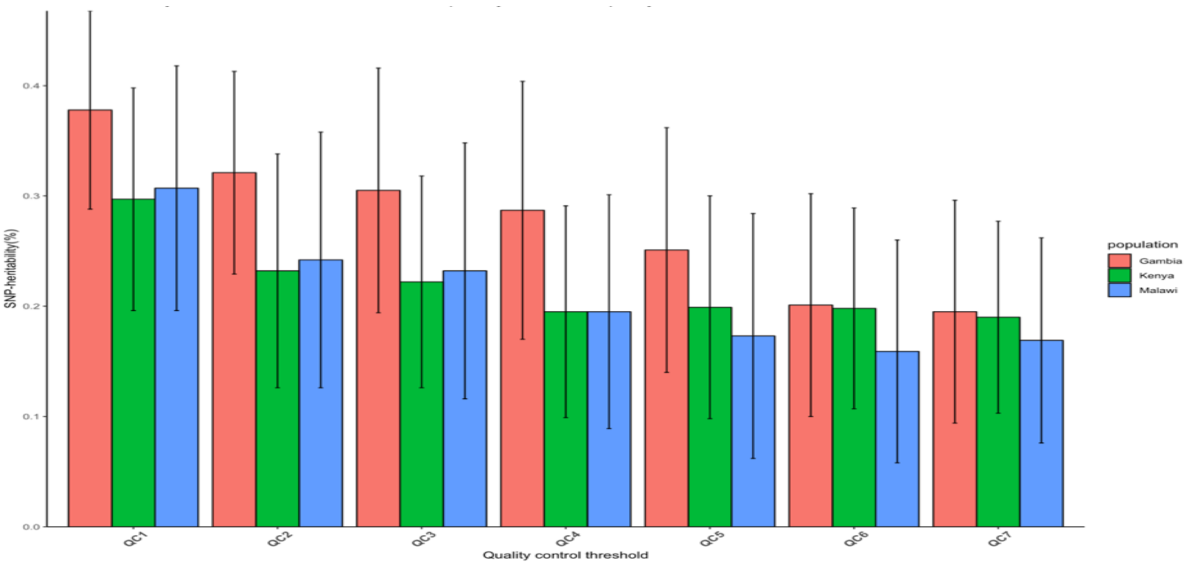


**Supplementary Figure 1**. SNP-heritability of severe malaria susceptibility and resistance at different QC-threshold from less stringent to more stringent (from left to right) determined by GCTA. Error bars represent the 95% CI of the estimates. QC1: GWAS QC-filtered data, QC2:QC1+ samples with 5% relatedness removed, QC3: QC2+ SNPs with differential missing rates $1x10^{-3}$ removed, QC4:QC3+ SNPs with missingness proportion $2 \times 10^{-2}$ removed, QC5: Q4+15 PCs as covariate, QC6: QC4+20 PCs, QC7=QC4+50PCs.

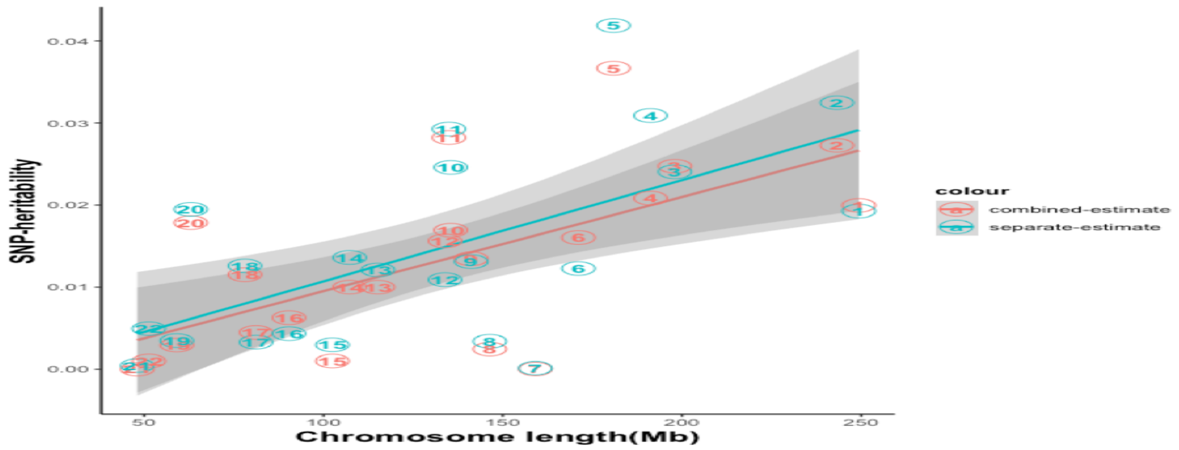


**Supplementary Figure 2.** SNP-heritability estimate per chromosome(y-axis) plotted against chromosome length(x-axis). The red line and blue line represent regressed SNP-heritability estimates obtained from joint GREML and separate analysis respectively. The 95% confidence interval around the slope of the regression model is represented by the grey shaded areas.

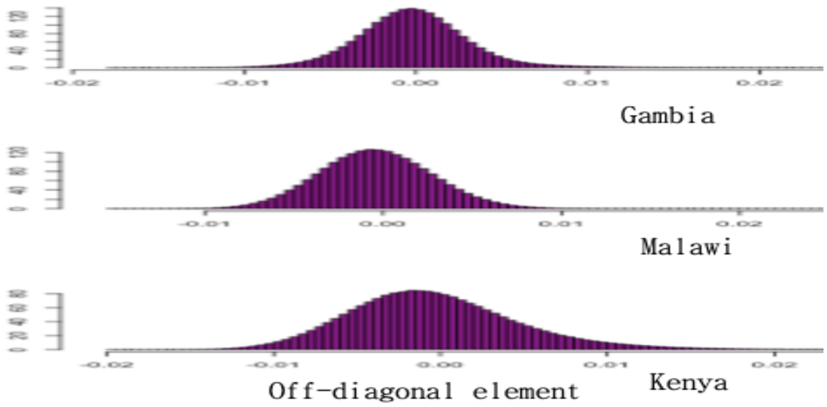


**Supplementary Figure 3.** Distribution of off-diagonal elements of the Genetic Relatedness Matrix (GRM) after removing closely related individuals (pairwise relatedness threshold >5%). As expected, the distribution was centred at zero suggesting that there was no close relatedness in all the datasets.

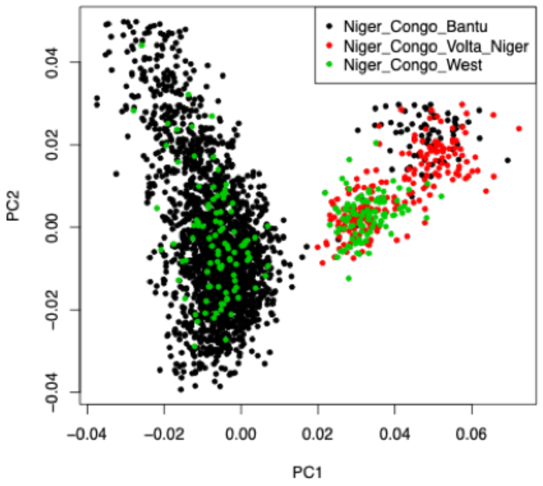


**Supplementary Figure 4.** We clustered our reference panel (quality filtered) in to geographic sub-regions in Africa using smart pca software. The left cluster represents east African population panel composed of 2112 samples and 22,473,268 SNPs. The right cluster represents the west African population panel composed of 380 samples and 18,919,068 SNPs.
